# Competing differentiation gradients coordinate fruit morphogenesis

**DOI:** 10.1101/2023.01.19.524793

**Authors:** A. Gómez-Felipe, M. Marconi, E. Branchini, B. Wang, H. Bertrand-Rakusova, T. Stan, J. Burkiewicz, S. de Folter, A-L. Routier-Kierzkowska, K. Wabnik, D. Kierzkowski

## Abstract

Morphogenesis requires the coordination of cellular behaviors along developmental axes^1^. In plants, gradients of growth and differentiation are typically established along a single longitudinal primordium axis to control organ shaping^2^. Here we combine quantitative live-imaging at cellular resolution with genetics, chemical treatments, and modeling to understand the formation of *Arabidopsis thaliana* female reproductive organ (gynoecium). We show that, contrary to other aerial organs, gynoecium shape is determined by two competing differentiation gradients positioned along two orthogonal axes. An early mediolateral gradient, dependent on meristematic activity in the medial domain, controls the valve morphogenesis while simultaneously restricting an auxin-dependent, longitudinal gradient to the style. This gradient competition serves to finetune the common developmental program governing organ morphogenesis to ensure the specialized function of the gynoecium^3,4^.

## MAIN TEXT

Morphogens provide positional information to control growth and patterning during the development of multicellular organisms^5,6^. Coordination between developmental gradients across different growth axes is critical for the establishment of a proper body plan during early embryogenesis and for the development of planar organs both in plants and animals^7,8^. In aerial organs of plants such as leaves, petals, or sepals, gradients of growth and differentiation are typically established along the longitudinal axis of the primordium to control their shape^2,9-11^. These gradients are proposed to be controlled by global organizers or biomechanical forces that coordinate cellular behaviors at the organ level^10-14^. However, what mechanisms and molecular components could account for global organizers remains unclear. Additionally, new growth axes can also be introduced locally through gene and hormone activities at the organ margins to produce protrusions such as leaflets or serrations in the leaf^15,16^.

Gynoecium, the female reproductive organ of the flower, is composed in *Arabidopsis thaliana* of two carpels fused with the repla and topped with the style and stigma (Fig. 1c)^3,4^. Gynoecium morphogenesis is suggested to be governed by a global polarity field spanning the epidermis and oriented parallel to the organ’s longitudinal axis^17^. However, in contrast to other aerial organs, longitudinal gradients of growth and differentiation have not been previously reported in *A. thaliana* gynoecium^17,18^. This remarkable exception is surprising as carpels have been shown to have a leaf-like origin^19^. It raises the question of how a common developmental program, likely governing organ shaping in plants^16,20-22^, could be modified to control the gynoecium morphogenesis to ensure its specialized function.

**Fig. 1.**
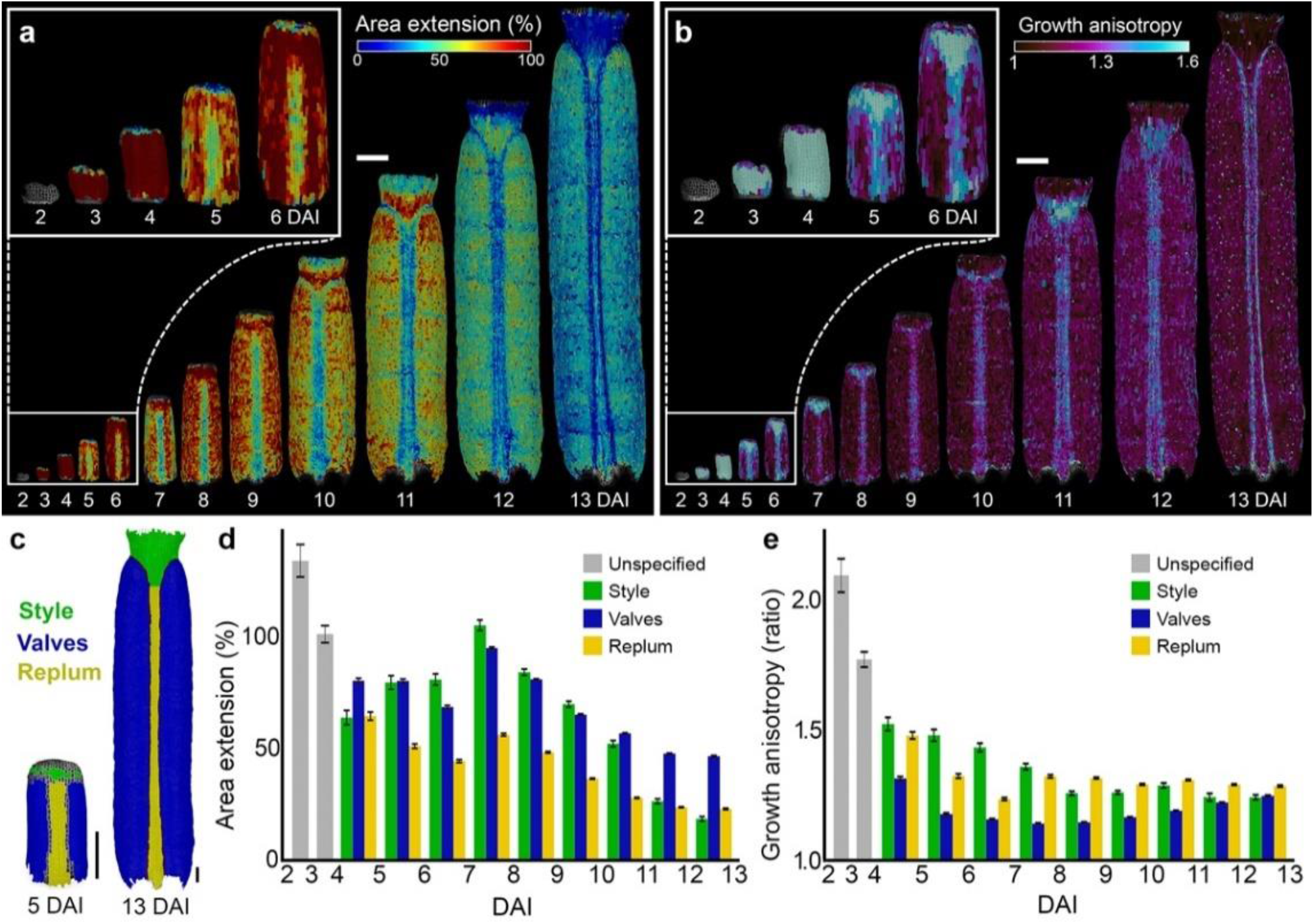
Cellular growth patterns underlying gynoecium development. **a-b**, Heat-maps of area extension(a) and growth anisotropy (b) for the *Arabidopsis thaliana* gynoecium. **c**, Lineage tracing of style (green), valves (blue) and replum (yellow) between 5 and 13 DAI. **d-e**, Quantifications of area extension (d) and growth anisotropy (e) in different regions of the developing gynoecium (Three independent time lapse series, n=104-10604 cells in one sample at specific time point). Error bars indicate SE. DAI: days after gynoecium initiation. Scale bars, 100 µm. See also Extended Data Fig. 1.

Gynoecium develops inside the enclosed floral bud, and it is very challenging to image and measure its growth, thus hindering our understanding of underlying processes. To overcome this limitation, we developed a new imaging method to precisely follow the cellular dynamics guiding gynoecium formation in *A. thaliana* (see Methods). We recorded the cellular behavior from early stages at 2 days after gynoecium initiation (2 DAI, equivalent of floral stage 7) until its final shape was established at 13 DAI^23^. Using MorphoGraphX software^24^, we extracted growth dynamics for all cells in the surface layer of one side of the gynoecium (from ∼70 cells at early primordium to ∼11000 cells at anthesis) and quantified growth rates, growth anisotropy (the ratio of expansion in the maximal and minimal principal directions of growth), cell divisions, and cell geometries (i.e., cell sizes) (Fig. 1a-b, and Extended Data Fig. 1a-b). We used lineage tracing^16^ to determine the origin of main gynoecium regions (i.e., valves, style, replum) and quantified their cellular growth parameters (Fig. 1c-e and Extended Data Fig. 1c-d).

In the early stages (from 2 to 4 DAI), cellular growth and proliferation were high, and the gynoecium expanded along its longitudinal axis (Fig. 1a-b,d-e, and Extended Data Fig. 1a,c). Between 4 and 5 DAI, regions that will develop into valves, replum, and style started to display divergent growth behaviors. The future style continued to elongate rapidly, while cell growth in the replum strongly decreased but maintained its longitudinal orientation. Valves continued to grow relatively fast but transitioned to much more isotropic growth (Fig. 1a-e). Consistent with previous studies^17,18^, cell area extension within both replum and valves appeared to be relatively uniform. By contrast, we observed a non-homogeneous decrease of growth in the style (from 9 DAI), with cells located at the tip of the style slowing down their growth earlier than at the base (Fig. 1a). These findings indicate that the common basipetal (from the tip to the base) gradient of growth observed in all *Arabidopsis* aerial organs is also present in the gynoecium, but it is established much later during development (around 8 DAI) and is spatially-restricted to the style.

This restriction suggests that cellular growth may arise from local, temporal gradient activities coordinated independently across different parts of the developing gynoecium.

To verify this scenario, we next computed cellular growth rates in different regions of the gynoecium along its longitudinal and mediolateral axes. The longitudinal growth was largely homogeneous in the valves, while we observed a clear basipetal (from the tip to the base) gradient of growth in the style from 7 DAI (Fig. 2a,c,d-e). Interestingly, upon valve specification at 4 DAI, we noticed an unequal cellular growth along the mediolateral axis of the organ (Fig. 2b-c). In particular, valve cells located close to the replum strongly increased their growth, establishing a clear gradient of cellular expansion along the mediolateral axis (Fig. 2b-c). This gradient persisted until 7/8 DAI when growth along the mediolateral axis equalized in the valves (Fig. 2b-c, and Extended Data Fig. 2b). Interestingly, we also observed corresponding gradients of cellular differentiation revealed by the appearance of the specialized cells (stomata or papilla) occurring first in the more distal regions of the style and more lateral regions of the valves (Fig. 2f-g, and Extended Data Fig. 1b). Our data suggest that time-shifted gradients of cellular behaviors are controlled independently along longitudinal and mediolateral axes in different parts of the developing gynoecium. Thus, two developmental gradients could locally tailor gynoecium morphogenesis along these perpendicular growth axes at different developmental windows.

**Fig. 2.**
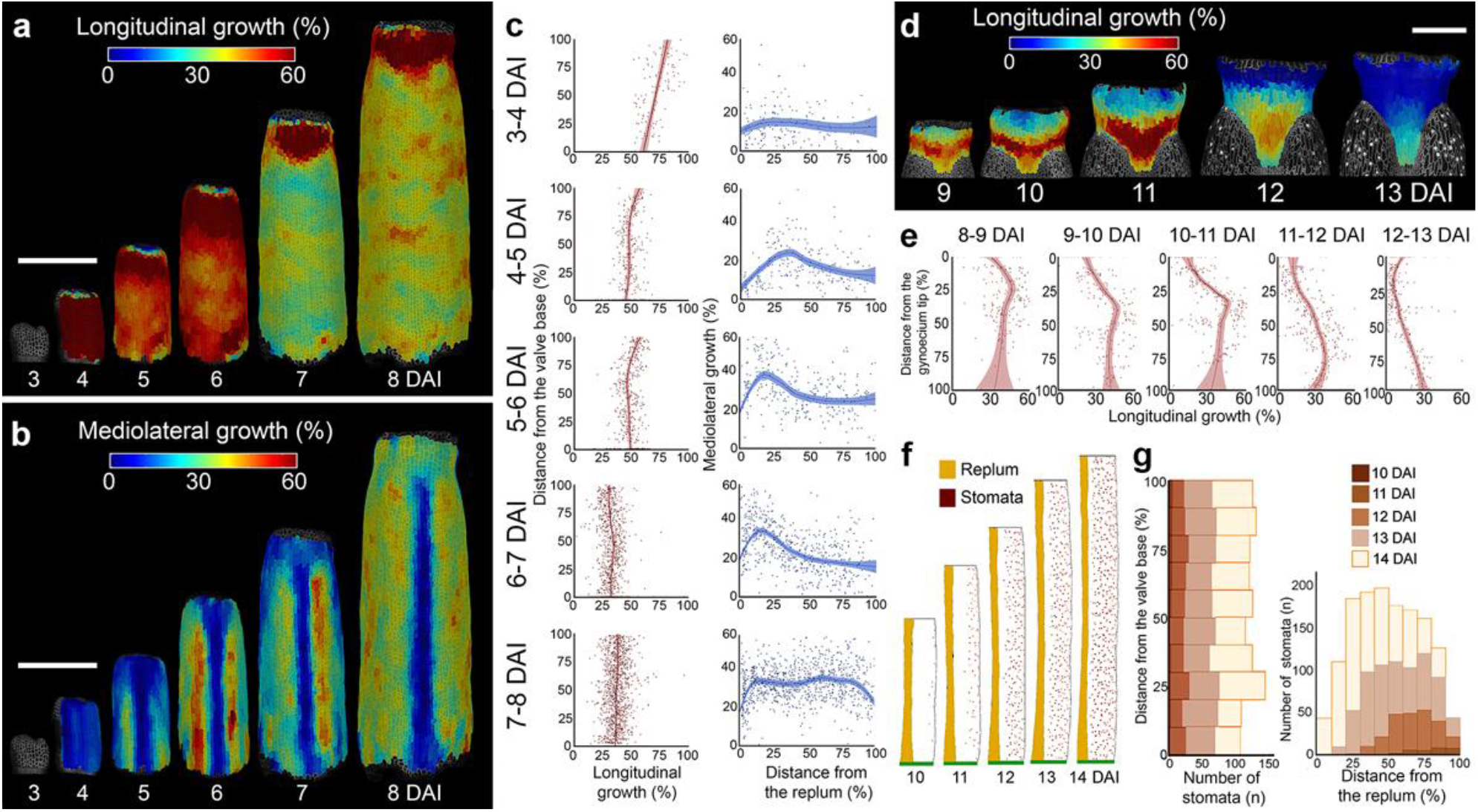
Two perpendicular gradients of growth and cell differentiation occur during gynoecium development. **a-b**, Heat-maps of the averaged cellular growth along longitudinal (a) and mediolateral (b) axes of the gynoecium. **c**, Quantification of the cellular growth along the longitudinal (left) and mediolateral (right) axis of the valve as a function of the distance from the valve base or from the replum (n=104-943 cells). **d**, Heat-maps of averaged cellular growth along the longitudinal axis of the style. **e**, Quantification of the cellular growth along the longitudinal axis of the style as a function of the distance from the gynoecium tip (n=207-321 cells). **f**, Stomata distribution in the valves. Replum in orange, valve in white, stomata in brown, base of the valve in green. **g**, Quantification of stomatal distribution as a function of the distance from the valve base (left) or from the replum (right) (n=6-2941 stomata). DAI: days after gynoecium initiation. For scatter plots, the distance was normalized. Dots in (c,e) indicate each growth value, lines represent the average and shaded areas represent standard deviation (SD). Scale bars, 100 µm in (a-b) and 50 µm in (d). See also Extended Data Fig. 2.

The molecular cues that could set these gradients are largely unknown. In theory, a longitudinal gradient may emerge from a hypothetical signal diffusing from the organ base that would prevent cellular differentiation in more proximal regions of the organ^12,16^. However, this scenario is unlikely in the style as the establishment of gradients in this part occurs at late developmental stages (from 8-9 DAI) when the style is located far away from the base of the organ. Alternatively, the longitudinal gradient could be controlled by a small molecule produced at the organ tip that would stimulate cell differentiation. The phytohormone auxin is a plausible candidate for such a signal, as auxin is known to control both cell growth and differentiation through concentration gradients in plants^16,25-27^. Auxin was also shown to control gynoecium patterning along its main axis^28^.

To test this hypothesis, we first monitored auxin responsiveness in the gynoecium using the DR5v2::nls-3xVenus reporter line^29^. DR5v2 signal in the epidermis was localized at the very tip of the gynoecium until anthesis, with its intensity decreasing progressively at late developmental stages from around 7-8 DAI (Fig. 3a and Extended Data Fig. 3a). Auxin synthesis monitored with YUCCA4 (pYUC4::YUC4:YFP) reporter line^30^ was initially detected in all epidermal cells (at 2 DAI), then became restricted to the replum (3-5 DAI) and to the very tip of the style (3-10 DAI) and was finally eliminated from the epidermis at around 11 DAI (Fig. 3c, and Extended Data Fig. 3b). The PIN-FORMED1 (PIN1) auxin efflux carrier was present at the upper membranes in all epidermal cells of the gynoecium at early stages (2-3 DAI) and later was progressively restricted to the replum and the style, consistent with its involvement in the apical accumulation of auxin^31^ (Fig. 3e-g). Interestingly, the first signs of the establishment of the basipetal gradient of growth in the style (from 8 DAI) coincided with the elimination of the PIN1 expression in the epidermis (Fig. 1a, Fig. 2d; Extended Data Fig. 3c). This suggests that acropetal (from the base to the tip) auxin transport in the style could help restrict auxin to its tip at early developmental stages to prevent precocious epidermal cell differentiation. At later stages, when PIN1 is eliminated from the epidermis, auxin could move through other PINs to more proximal regions triggering the basipetal gradient of cell differentiation.

**Fig. 3.**
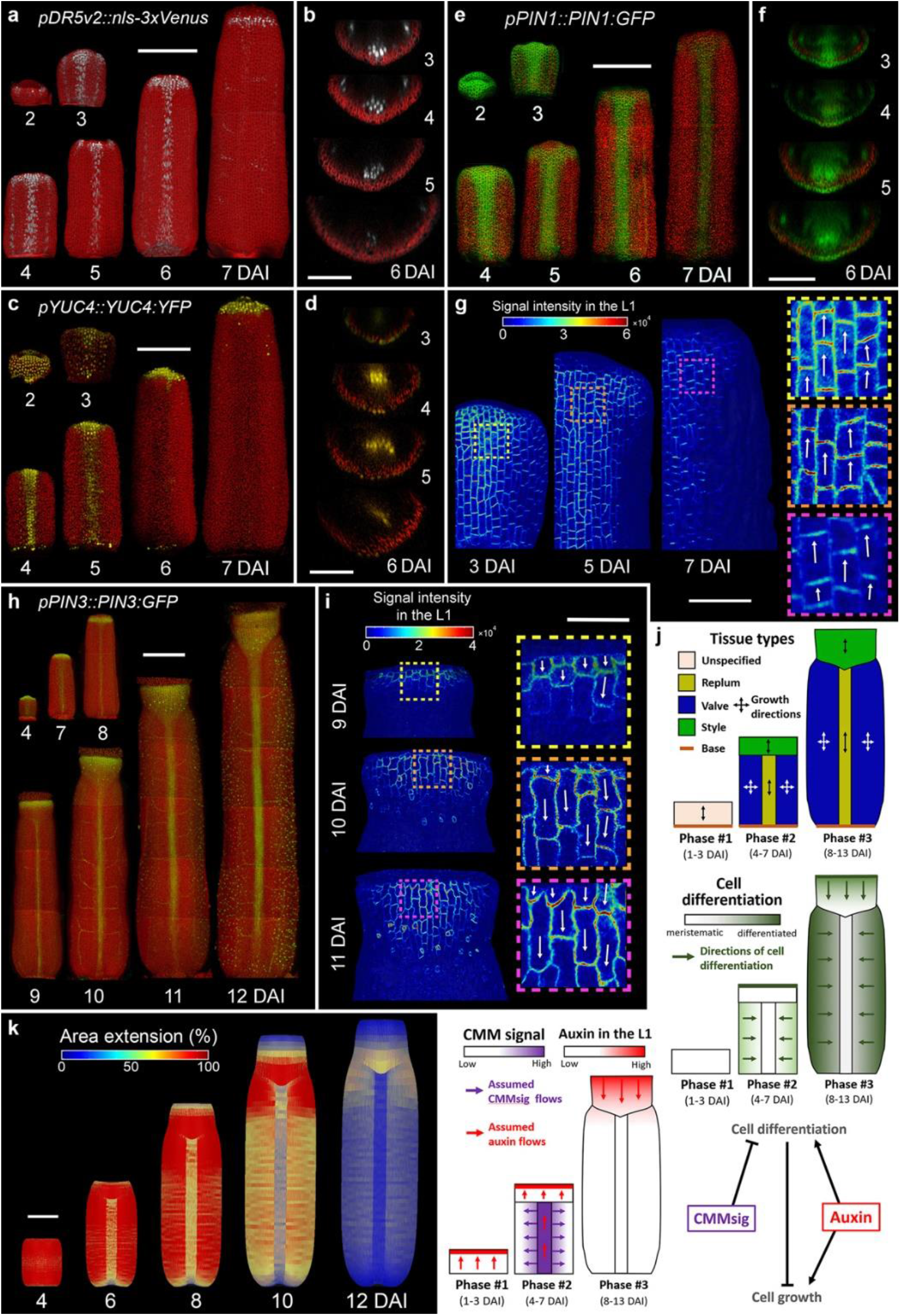
Auxin patterning and the model of gynoecium development. **a–k**, Expression patterns of *pDR5v2::nls-3xVenus* (a-b), *pYUC4::YUC4:YFP* (c-d), *pPIN1::PIN1:GFP* (e-g), and *pPIN3::PIN3:GFP* (h-i) in the *A. thaliana* gynoecium. Virtual cross sections shown in (b, d, f) are located in the middle of the gynoecium. **g, i**, Heat-maps represent the intensity of the *PIN1:GFP* (g) and *PIN3:GFP* (i) signal in the L1 epidermal layer (one side of the gynoecium tops are shown for PIN1 and the style is shown for PIN3). Insets indicate membrane localization and arrows indicate orientation of PIN polarization. **j**, Model of *A. thaliana* wild-type gynoecium growth. **k**, Resultant shape and distribution of growth rates predicted by the model for wild-type gynoecium development. DAI: days after gynoecium initiation. Scale bars, 100 µm in (a, c, e, h, k) and 50 µm in (b, d, f, g, i). See also Extended Data Figs. 3-5.

To explore this possibility, we compared the expression of other PIN proteins involved in gynoecium patterning with growth and differentiation gradients in the style. PIN3 was initially (until 8 DAI) expressed at the tip of the style in a non-polarized manner consistent with its role in style radialization^31,32^ (Fig. 3h). Strikingly, from 9 DAI, the PIN3 expression domain started to extend toward more proximal regions of the style, tightly following the gradient of cellular growth and differentiation in this region (Fig. 3h-i). Importantly, PIN3 appeared to switch its polarization away from the organ tip (Fig. 3i), suggesting progressive establishment of basipetal auxin flow through the style at late developmental stages. Such basal polarization was previously observed only in the epidermis of the ovary after fertilization^33^. Consistently, PIN7 followed a comparable pattern of distribution and polarization in the style epidermis from 9 DAI (Extended Data Fig. 3d-e). These data indicate that auxin originating from the gynoecium tip sets up the longitudinal gradient of growth and differentiation in the style at late developmental stages.

Why is this longitudinal gradient restricted to the style? During early gynoecium development, a carpel margin meristem (CMM) is formed along the margins at the inner side of the fused carpels. The CMM gives rise to the placentae which produce ovules^34,35^ and is critical for the establishment of the valves^32,36^. The activity of CMM monitored by the ovule primordia initiation and associated auxin production (YUC4 expression), auxin response (DR5v2), and PIN1-mediated auxin transport, coincided with the establishment of the mediolateral gradient of growth and differentiation in the valves from around 4-7 DAI (Fig. 2b; Fig. 3a,c,e; Extended Data Fig. 4)^37^.

Therefore, we hypothesize that an active CMM may be a source of a putative mobile signal that sets the mediolateral gradient of differentiation at early developmental stages of the gynoecium. Furthermore, as the mediolateral gradient is the first to be set during gynoecium development, we propose that it eventually outcompetes the latter longitudinal gradient by desensitizing valves to a tip-derived signal by triggering valve cell differentiation at early stages.

To test this theory, we constructed a spatial-temporal computer model of the gynoecium that integrates our experimentally-derived assumptions (Fig. 3j, see Methods). Briefly, the simulation follows three stages representing the most relevant phases of gynoecium development (Fig. 3j). Phase#1 (1-3 DAI): early gynoecium is composed of undifferentiated cells, and it grows rapidly along the longitudinal organ axis. Phase#2 (4-7 DAI): new gynoecium regions (valves, replum, and style) emerge, displaying divergent growth dynamics (fast and predominantly isotropic growth in the valves, slow and anisotropic growth in replum, fast and anisotropic growth in the style). Phase#3 (7-12 DAI): the longitudinal gradient of cell growth and differentiation is set in the style and the organ achieves its final shape (Fig. 3j). The core assumptions of the model are: (1) non-differentiated cells grow along the longitudinal axis of the organ; (2) Once the replum is established (Phase#2), its growth is reduced and restricted to the longitudinal axis (Fig. 1a and Fig. 2a,b); (3) CMM is the source of a putative mobile signal (*CMMsig*) that triggers the identity of the valve and inhibits differentiation in a concentration-dependent manner (Fig. 2f); (4) Auxin moving through the epidermis leads to the establishment of a growth and differentiation gradient in the style during Phase#3 (Fig. 3j). Valves show less competence to respond to tip derived auxin signals during Phase#3 as they already started differentiating along the mediolateral axis during Phase#2. With these assumptions, the computer model recapitulated general patterns of gynoecium growth that were comparable to that observed experimentally (Fig. 1a, Fig. 3k, and Extended Data Fig. 5).

Similarly to experimental data, mediolateral growth concentrated solely in the valves and the longitudinal gradient of growth was restricted to the style (Extended data Fig. 5a-b).

Our model assumes that the basipetal gradient set by auxin at later developmental stages (from 7-8 DAI, Phase#3) is mainly restricted to the style by the activity of the CMM which controls the establishment of the mediolateral gradient of cell differentiation in the valves during Phase#2, making them less competent to respond to basipetal signal during Phase#3. Thus, removing the CMM should allow the longitudinal gradient to spread to the more proximal regions of the organ as they should be composed of undifferentiated cells that are now competent to respond to auxin coming from the organ tip. Eliminating the CMM activity in our simulations abolished the establishment of the valves resulting in the formation of radially symmetric organ instead of the ovary (Fig. 4a, Extended Data Fig. 6). Importantly, since the CMM is now missing, the regions that correspond to valves in wild-type scenarios are now competent to respond to the basipetal signal resulting in expansion of gradients of cellular growth and differentiation (Figs. 3k and 4a, and Extended Data Fig. 6).

**Fig. 4.**
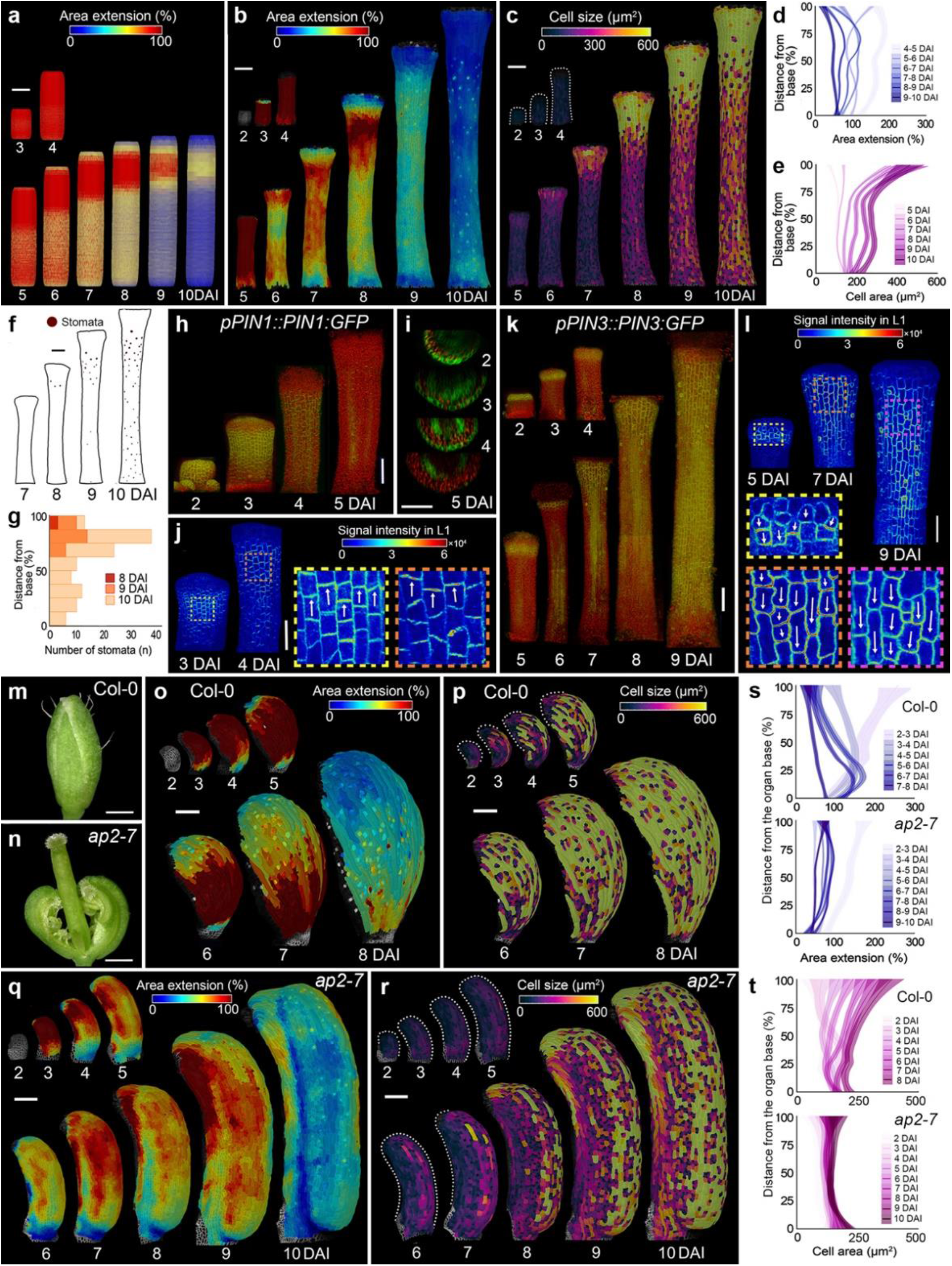
Marginal meristem activity underlies the establishment of the mediolateral developmental gradient. **a**, Resultant shape and distribution of growth rates predicted by the model of the gynoecium without the CMM. **b-c**, Heat maps of averaged area extension (b) and cell sizes (c) in *A. thaliana* gynoecia treated with the NPA. **d-e**, Quantification of cellular growth (d) and cell sizes (e) as a function of the distance from the gynoecium base after NPA treatment (three independent time-lapse series, n=219-1479 cells). **f-g**, Stomata distribution (f) and quantification of stomatal distribution as a function of the distance from the gynoecium base (g) after NPA treatment (three independent time lapse series, n=3-100 stomata). **h-l**, Expression patterns of *PIN1:GFP* (h-j) and *PIN3:GFP* (k-l) in gynoecia after NPA treatment. Heat maps represent the intensity of the *PIN1:GFP* (j) and *PIN3:GFP* (l) signals in the L1 layer. Insets indicate membrane localization; arrows indicate the orientation of PIN polarization. **m-n**, Wild-type (m) and *ap2-7* sepal (n). **o-r**, Heat maps of averaged area extension (o,q) and cell sizes (p,r) of wild-type (o-p) and *ap2-7* (q-r) sepals. **s-t**, Quantifications of area extension (s) and cell sizes (t) of wild-type sepal (three independent time lapse series, n=113-1142 cells) and *ap2-7* sepal (three independent time lapse series, n=44-1478 cells). For plots, the distance was normalized, lines represent the average and shaded areas represent standard deviation (SD). DAI: days after organ initiation. Scale bars, 100 µm (b-c, k-r) and 50µm (i-j, l). See also Extended Data Figs. 5-9.

To experimentally test this model prediction, we removed the CMM activity using naphthylphthalamic acid (NPA) treatment during gynoecium initiation. Indeed, this chemical treatment led to the complete elimination of the valves and conversion of the gynoecium into a radially symmetric tube-like structure^28,36^, similar to that observed in model simulations (Fig. 4a-c, and Extended Data Fig. 6). In this valveless gynoecium, the mediolateral growth was abolished while the organ mainly expanded longitudinally (Extended Data Fig. 7). As in the wild-type, we observed an establishment of the longitudinal gradient of growth and differentiation in the region corresponding to style at later developmental stages (from 7 DAI). However, those gradients were extended farther toward more proximal regions of the organ as compared to the wild-type where they were mainly restricted to the very tip of the gynoecium (Figs. 1a, 4b-g, and Extended Data Fig. 5a). The onset of the longitudinal gradients happened after the complete elimination of the apically polarized PIN1 auxin efflux carrier from the epidermis (Fig. 4h-j, and Extended Data Fig. 6a). Strikingly, PIN3 that was initially (until 5 DAI) localized at the tip of the gynoecium in a non-polarized manner, started (from 5-6 DAI) to polarize basally and progressively expanded its expression throughout the entire epidermis at 9 DAI (Fig. 4k-l). This suggests that auxin, which was initially synthesized and accumulated at the gynoecium tip (Extended Data Fig. 8b-d), was likely transported through the epidermis toward the organ base to trigger the differentiation along the longitudinal axis of the gynoecium. These results support the idea that an early CMM-dependent establishment of the mediolateral gradient of differentiation in the valves is key to preventing the tip-derived signal from setting up an organ-wide basipetal gradient of cell differentiation.

If the presence of the CMM prevents cell differentiation from progressing along the main axis of the gynoecium, the introduction of ectopic meristematic activity along the margins of other lateral organs should abolish their typical basipetal gradients of growth and differentiation^12,16,36,38^. In the wild-type sepals (Fig. 4m), a clear longitudinal gradient of cellular growth and differentiation was visible from around 5 DAI (Fig. 4o-p,s-t, and Extended Data Fig. 9a-c)^11^. We then analyzed the growth in the *ap2-7* mutant sepals, which develop carpeloid structures including placenta and ovules along their margins (Fig. 4n,q)^39^. In such carpelized sepals, the typical basipetal gradient of cellular growth and differentiation was largely eliminated (Fig. 4q-t and Extended Data Fig. 9d-f). Furthermore, cells located close to the modified margin of the *ap2-7* sepal tended to differentiate later than cells further away from the margin, indicating that the meristematic activity along the carpelized sepals in this mutant led to the establishment of the mediolateral gradient of differentiation (Fig. 4r and Extended Data Fig. 9f). These findings confirm that the CMM activity prevents the progression of basipetal cell differentiation throughout the entire organ by setting up the mediolateral gradient early during development.

Our study revealed an underlying mechanism shaping gynoecium development, alternative to that of other aerial organs. It is directed by competition between orthogonal time-shifted differentiation gradients. A general delay of cell differentiation in the gynoecium is one of the critical events for proper patterning of the fruit. It allows the introduction of the mediolateral gradient in the valves before the onset of the basipetal cell differentiation. The mediolateral gradient in turn restricts the progression of the basipetal gradient, typical for all lateral organs, to the style. From the evolutionary point of view, this novel feature has contributed to the success of flowering plants enabling simultaneous maturation of ovules that is critical for successful plant reproduction. Also, we identified that two plausible signals control orthogonal gradients of differentiation, one likely being auxin itself^28,31-33^. The second signal could be other type of small molecules that are known to act in a concentration-dependent manner including plant hormones^40,41^. Cytokinin is particularly plausible candidate for such a signal as it is active in the CMM, can diffuse through the tissue, and is known to delay cell differentiation^32,34,36,42^. Finally, the role of the mediolateral signaling is also supported by recent study showing that putative signal derived from the developing seeds is critical for fruit expansion after fertilization^18^.

There are interesting similarities of proposed mechanism with morphogen competition mechanisms operating along single axis of growth in synthetic and natural systems^43,44^. However, unlike other studies this work highlights the importance of gradients that can act on different axes of growth but also in different time windows, thus providing new insights into mechanisms of spatial developmental competition during multicellular development. Our work will help expand the repertoire of principles and strategies for the controllable regulation of morphogenesis through spatial-temporal finetuning of morphogen gradients, thereby strengthening a foundation for applications in the emerging field of synthetic developmental biology^45,46^.

## METHODS

### Plant material and growth conditions

*pUBQ10::myb:YFP*^47^, *pUBQ10::PM:TDTomato*^48^, *pPIN1::PIN1:GFP*^49^, *pPIN3::PIN3:GFP*^50^, *pPIN7::PIN7:GFP*^51^, *pYUC4::YUC4:YFP*^30^, and *ap2-7*^39^ (Accession number N6241) were in Columbia background and *DR5v2::nls-3xVenus*^29^ was in Columbia/Utrecht background. *pUBQ10::PM:TDTomato* and *DR5v2::nls-3xVenus* were crossed and analyzed in F3. Plants were grown on soil in a growth chamber under long-day conditions (16 h illumination, 95 µmol m2 s1) with 60%–70% relative humidity at 22 ±1°C.

### Live-imaging microscopy

3-weeks-old flowers were dissected. Inflorescences were cut 2 cm below the apex, and the oldest floral buds were removed using fine tweezers to uncover young floral primordia. Floral buds at stage 7 and 8^23^ were selected and initiating sepals, petals and stamens were manually removed with a needle and fine tweezers to expose the early gynoecium. Dissected floral buds were transferred to Ø60 mm petri dishes with 1/2 Murashige and Skoog (MS)^52^ medium supplemented with vitamins, 1.5% agar (w/v), 1% sucrose (w/v), and 0.1% Plant Protective Medium (Plant Cell Technologies; v/v). Dissected stems were placed horizontally in a cavity cut out in the culture medium as described previously^53^, immersed in water, and abaxial gynoecium surface was imaged at 24 h intervals for up to 13 days. Between imaging, water was removed, and samples were transferred to a growth chamber with standard long day conditions (16 h illumination, 95 µmol m2 s1) with 60%–70% relative humidity at 22 ± 1°C. The images shown in Fig. 1 and 2 are derived from two independent overlapping time lapse experiments (2-4 DAI and 4-13 DAI). The images shown in Fig. 4b-c are derived from two independent overlapping time lapse experiments (2-5 DAI and 5-10 DAI). The images shown in Fig. 4q-r are derived from two independent overlapping time lapse experiments (2-4 DAI and 4-10 DAI).

### Microscopy and image analysis

All the confocal imaging was performed using a Zeiss LSM 800 confocal laser scanning upright microscope with a 40x long-distance working, water-dipping objective (W Plan-Apochromat 40x/1.0 DIC VIS-IRM27). Excitation was achieved using a diode laser with 488 nm for YFP and GFP, and 561 nm for TdTomato. The emission was collected at 500-550 nm for GFP, at 490-520 nm for YFP/Venus, and at 600-660 nm for TdTomato. Confocal stacks were acquired using a step size of 0.5-1 µm distance in *z*-dimension, at 16 bits image depth, and 512 × 512 pixel resolution. For samples that were larger than the microscope field of view, multiple overlapping stacks were acquired and stitched using MorphoGraphX^24,54^.

### Image analysis

The resulting confocal images were processed and analyzed using the MorphoGraphX software^24,54^. Stacks were manually stitched into a single file using the “combine stack” tool. Surface detection was performed with the “edge detect” tool with a threshold from 8,000–10,000 followed by “edge detect angle” (threshold 4,000–6,000). Initial meshes of 3-5 µm cube size were created and subdivided 2-3 times before projecting the membrane signal (2-4 µm from the mesh). Segmentation was performed with the “auto-segmentation” tool, followed by manual curation and segmentation of additional cells at the periphery of the mesh. Parent relations between cells of successive time points were attributed manually.

For lineage tracking analysis, the parent relations between each of the consecutive time points were combined to compute corresponding cell lineages over multiple days. The lineage data were later used to compute area extension, cell growth anisotropy, and cell proliferation. Area extension is calculated as the relative increase between the surface area of a mother cell and the area of its daughter cell(s) at the next time point. It was expressed in percentage increase ([Total area of daughter cells at T1/Area of mother cell T0-1] x 100). Heat maps of growth, growth anisotropy, and cell divisions generated between consecutive time points were displayed in the second time point.

Growth rates along the longitudinal and mediolateral axis of gynoecia were calculated using a custom Bezier grid that was manually placed so that it closely followed the geometry of the organ at each time point. They were expressed as percentage increase ([Total length of daughter cells at T1/length of mother cell T0-1] x 100). Distance from the filament base was calculated using the “Cell distance” tool, which measures the shortest path between cells following the organ surface geometry.

Reverse lineage tracing of cells corresponding to different organ regions (style, replum and valves) was performed as described previously^53^.

Heat-maps of membrane localization of PIN proteins (*PIN1:GFP, PIN3:GFP, PIN7:GFP*) were performed as described previously^16^. GFP signal was projected from 2-4 µm from the extracted mesh.

### Chemical treatment

For 1-N-naphthylphthalamic acid (NPA), flowers were treated for three consecutive days one week after bolting with the solution containing 30 μM 1-N-naphthylphthalamic acid (Sigma) and 0.010 % (v/v) Silwet L-77. All treated plants were then grown under long-day conditions (16 h illumination, 95 µmol m2 s1) with 60%–70% relative humidity at 22 ±1°C. One week after the last NPA treatment, flowers were dissected for live-imaging.

### Model description

The computational model simulations were implemented using MorphoDynamX, a modeling platform based on MorphoGraphX^24,54^. MorphoDynamX is an advanced modeling framework based on a geometric data structure called “Cell Complex”^55^. The model’s initial template was a triangulated hollow cylindrical mesh on which physical forces and chemical species are simulated. The gynoecium growth was implemented using Position Based Dynamics (PBD)^56^.

The model makes the following key assumptions:

1. Gynoecium development is divided into three phases: Phase#1: Early growth after initiation (1-3 DAI). The initial geometry of the primordium is a hollow tube of size 50 µm in length and 100 µm in width. The early gynoecium is assumed to be a homogenous tissue composed of undifferentiated cells. At this early stage, we assume that growth is strongly anisotropic and distributed uniformly along the organ. Phase #2: Establishment of clonally distinct regions (4-7 DAI). During this phase, we observe the appearance of regions that will clonally develop into style, replum, and valves (Fig. 1C). Based on our experimental observations, we assumed that cells in the valves start to differentiate and slowly acquire isotropic growth, while cells in the replum and style stay undifferentiated and conserve the initial longitudinal elongation. As a result, growth rates are reduced in the replum while style and valves grow fast. In the model, we assume that replum produces a mobile signal (*CMMsig*) promoting growth (by delaying differentiation) in the neighboring valve cells. *CMMsig* induces a gradient of growth in the valves as discussed later. Phase #3: The establishment of the longitudinal growth gradient in the style and the final shape acquisition (7-12 DAI). During this phase, we introduced the basipetal gradient of growth in the style that corresponds to the assumed auxin gradient generated by the activity of PIN3- and PIN7-driven basipetal auxin movement through the epidermis (*BPsig*). Valves show less competence to respond to this tip-derived auxin signal as their cells are already partially differentiated. During this phase, CMMsig is deactivated coinciding with the cessation of meristematic activity and ovules initiation. Finally, the components of the model stop growing, and the gynoecium reaches its final shape.
2. Global organ growth is driven by uniform internal pressure. This choice consents to replicate a realistic organ shape and at the same time to abstract from single-cell local growth. However, each triangle of the mesh is allowed to resist expansion depending on the concentration of the chemicals, as described later. The pressure acting on every single vertex of the mesh is expressed by the formula:

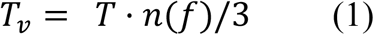

*Tv* is the force acting on vertex *v*; *Tv* is the turgor pressure force and *n*(f) is the normal vector to face *f*.
3. Local tissue growth is regulated through the modulation of the rest length of the mesh edges by chemical morphogens. The rest length is later used as a parameter for the distance constraint of the PBD engine. The rest length for a given edge is updated according to the formula:

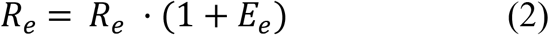

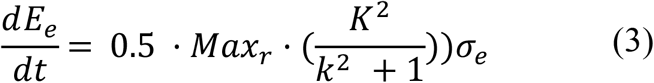

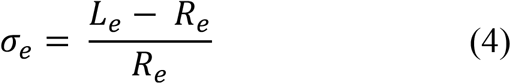

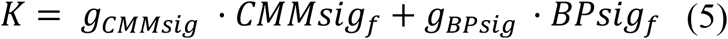

*Re* is the current rest length of edge *e*; *Ee* is the rest length update rate of edge *e*; *e* is the strain on edge *e*; *Maxr* is the maximum update rate; *K* is the total morphogens contribution on edge update rate; *Le* is the length of edge *e*; *CMMsigf* is the concentration of placenta signal inside face *f* (as described later); *BPsigf* is the concentration of basipetal signal inside face *f* (as described later); *gCMMsig* is the growth promotion coefficient of *CMM* signal; *gBPsig* is the growth promotion coefficient of the basipetal signal.
4. We assume that tissue growth is by default anisotropic in undifferentiated tissue. Local tissue anisotropy is represented using an anisotropy factor. This anisotropy factor is later used as a parameter for the strain constraint of the PBD engine (see below). The anisotropy factor ranges from zero (full isotropy) to 1 (full anisotropy). During stage 1, the whole organ is assumed to be anisotropic (as discussed before). At stage 2, valves start to acquire a more isotropic shape; this is achieved by relaxing the anisotropy factor:

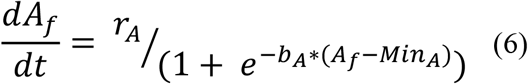

*Af* is the anisotropy factor for face *f*; *rA* is the anisotropy change rate; *MinA* is the half-minimum anisotropy factor; *bA* is the anisotropy factor change rate coefficient.
5. *CMMsig* is produced in the replum and diffuses in the surrounding tissues (mainly in the valves). In the valves, *CMMsig* creates a bell-shaped mediolateral gradient of growth. In the model, *CMMsig* diffusion is not directly modeled through morphogen exchange between neighboring faces as the high number of mesh elements would make this process computationally slow. Instead, we calculated the distance of each mesh faces from the morphogen source (replum) and calculated the morphogen concentration as a function of source expression rate and source distance:

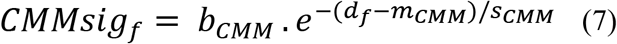

*CMMsigf* is the concentration of carpel margin meristem derived signal inside face *f; bCMM* is *CMMsig* maximum expression rate; *df* is the distance of face *f* from producer (the replum in this case); *mCMM* is the signal center of expression; *sCMM* is the signal surface spread.
6. A basipetal signal (*BPsig*) is generated along the style and is modeled in a similar fashion as *CMMsig* described before, with the source located at the very top of the organ. This signal appears later in the simulation (corresponding to 7-11 DAI, Fig. 1a). The signal generates a bell-shaped gradient centered in the style and induces tissue growth:

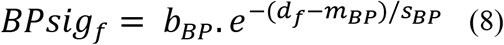

*BPsigf* is the concentration of basipetal signal inside face *f; bBP* is *BPsig* expression rate; *df* is the distance of face *f* from producer (the top of the organ in this case) *mBP* is the signal center of expression; *sBP* is the signal surface spread.
7. PBD is a modeling technique that overcomes the typical limitation of force-based systems (which tend to be unstable and/or computationally expensive) by omitting the velocity calculation layer and the working direction of vertex positions throughout the application of physical constraints. The current implementation of PBD is based on a more comprehensive adaptation applied for root growth^57^. The model presented here relies on two PBD constraints: the distance constraint^56^ which is applied to the mesh edges and regulates tissue stiffness and growth, and the strain constraint, which is applied to mesh faces and regulates tissue anisotropy^58^. The distance constraint is regulated by two parameters: the constraint stiffness for compression/stretching (*kc / ke*; Table 1) and the rest length of the edge (described in equation 2). The strain constraint is regulated by the anisotropy factor (described in equation 5), which serves as vertical constraint stiffness.

**Table 1.**
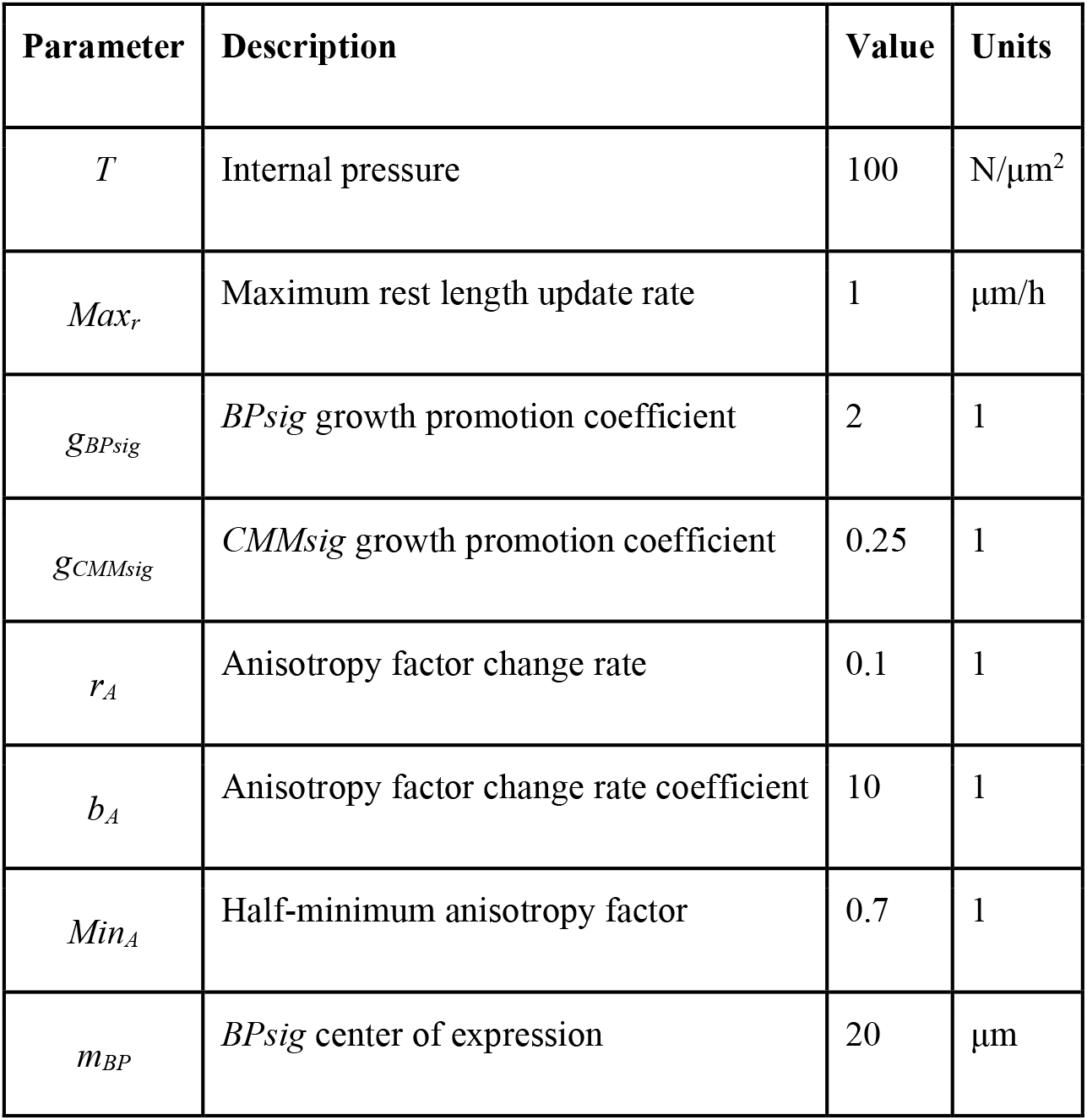
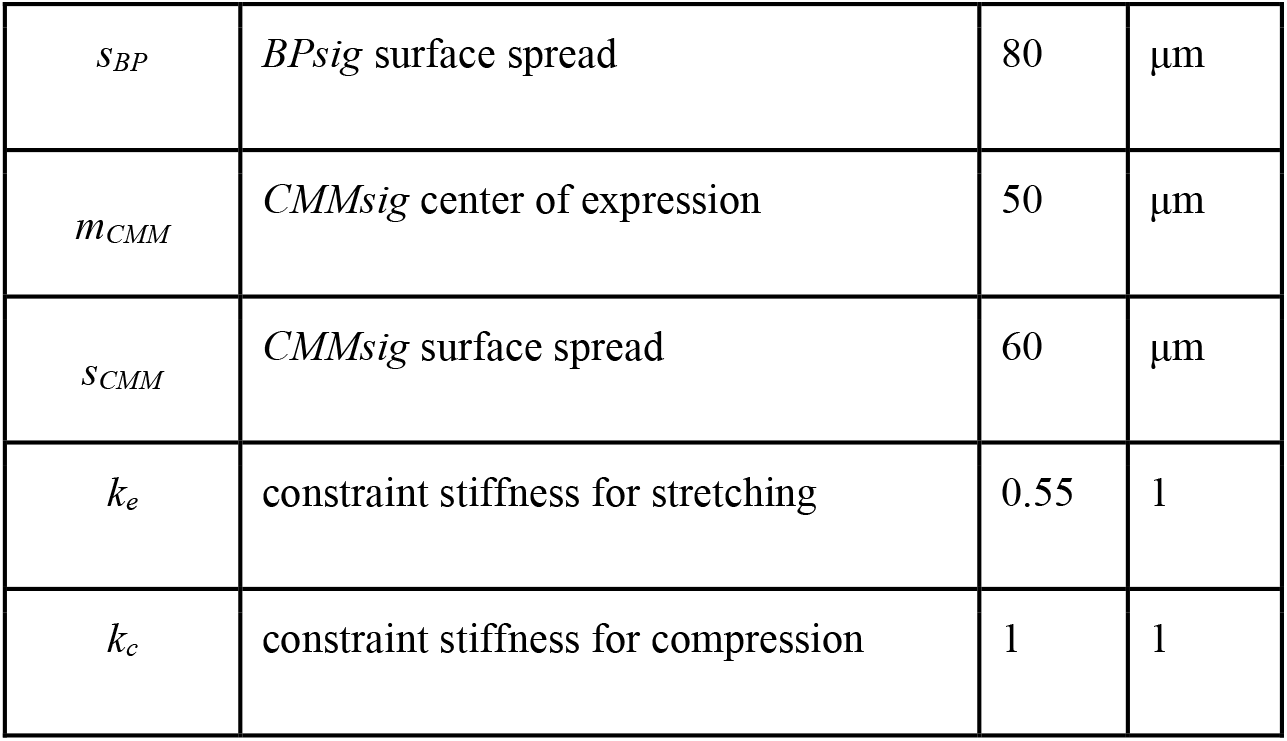
Model Parameters.
8. To replicate NPA treatment (Fig. 4b-c) *CMMsig* is abolished from the model, while replum and valves do not acquire their identity. Hence, the gynoecium maintains the initial anisotropic growth for the entirety of the simulation.

| Parameter | Description | Value | Units |
| --- | --- | --- | --- |
| $T$ | Internal pressure | 100 | N/$\mu\text{m}^2$ |
| $Max_r$ | Maximum rest length update rate | 1 | $\mu\text{m}/\text{h}$ |
| $g_{BPsig}$ | *BPsig* growth promotion coefficient | 2 | 1 |
| $g_{CMMsig}$ | *CMMsig* growth promotion coefficient | 0.25 | 1 |
| $r_A$ | Anisotropy factor change rate | 0.1 | 1 |
| $b_A$ | Anisotropy factor change rate coefficient | 10 | 1 |
| $Min_A$ | Half-minimum anisotropy factor | 0.7 | 1 |
| $m_{BP}$ | *BPsig* center of expression | 20 | $\mu\text{m}$ |
| $s_{BP}$ | $BP_{sig}$ surface spread | 80 | $\mu\text{m}$ |
| $m_{CMM}$ | $CMM_{sig}$ center of expression | 50 | $\mu\text{m}$ |
| $s_{CMM}$ | $CMM_{sig}$ surface spread | 60 | $\mu\text{m}$ |
| $k_e$ | constraint stiffness for stretching | 0.55 | 1 |
| $k_c$ | constraint stiffness for compression | 1 | 1 |

### Data and code availability statement

The confocal stacks and meshes used to extract growth parameters, all R-scripts used to analyze data and the model codes are available to download from the Open Science Framework repository (link to be provided).

## ACKNOWLEDGEMENTS

We thank Olivier Hamant and Lars Østergaard for the critical reading of the manuscript. We thank Carlos Montoya for help with the sample segmentations, and Dolf Weijers, Zhengjuan Zhang, and Kristoffer Jonsson for seeds of reporter lines. This work was supported by a New Initiative grant from Centre SEVE to D.K. and Fonds de Recherche du Québec Nature et Technologies Team Grant (2021-PR-282285) to D.K and A-L.R-K. D.K. was funded by Discovery grants from the Natural Sciences and Engineering Research Council of Canada (RGPIN-2018-05762). A-L.R-K. was funded by a Discovery grants from the Natural Sciences and Engineering Research Council of Canada (RGPIN-2018-04897). E.B. was funded by the Natural Sciences and Engineering Research Council of Canada graduate study grant (BRPC-549453-2020). K.W. was funded by The Programa de Atracción de Talento 2017 (Comunidad de Madrid, 2017-T1/BIO-5654) and PID2021-122158NB-I00 (2022-2025). In the frame of SEV-2016-0672 and CEX2020-000999-S funding to CBGP, M.M. was supported with a postdoctoral contract. K.W. was supported by Programa Estatal de Generación del Conocimiento y Fortalecimiento Cientifico y Tecnológico del Sistema de I+D+I 2019 (PGC2018-093387-A-I00) and I+D+I 2021 (PID2021-122158NB-I00) from El Ministerio de Ciencia, Innovación y Universidades (MICIU) (to K.W.). This work has been performed in the frame of the initiative “Centre of Excellence for Plant-Environment Interactions (CEPEI)” between the CBGP, UPM-INIA/The Spanish National Research Council (CSIC) and the IGDB and the Plant Stress Centre of the CAS. CEPEI initiative has been financially supported by the “Severo Ochoa Programme for Centres of Excellence in R&D” from the Agencia Estatal de Investigación of Spain [grants SEV-2016-0672 (2017 to 2021) to the CBGP]. S.D.F. was supported by the Mexican National Council of Science and Technology (CONACyT) grant CB-2017-2018-A1-S-10126.

## AUTHOR CONTRIBUTIONS

D.K. and A.G.F., conceived and designed the experiments; K.W. and M.M. conceived and designed the computational models; A.G.F. and H.B.R. performed all experiments with the help from B.W.; A.G.F., E.B., B.W., T.S., and J.B. extracted data from time-lapse series with the help from A-L.R-K.; A.G.F., B.W., and E.B. analyzed the data; D.K. supervised the project; A.G.F., M.M., K.W., and D.K., wrote the manuscript with input from S.D.F. and A-L-R-K; D.K., A-L.R-K., and K.W. provided the funding. all the authors reviewed the manuscript.

## COMPETING INTERESTS

The authors declare no competing or financial interests.

## EXTENDED DATA

**Extended Data Fig. 1.**
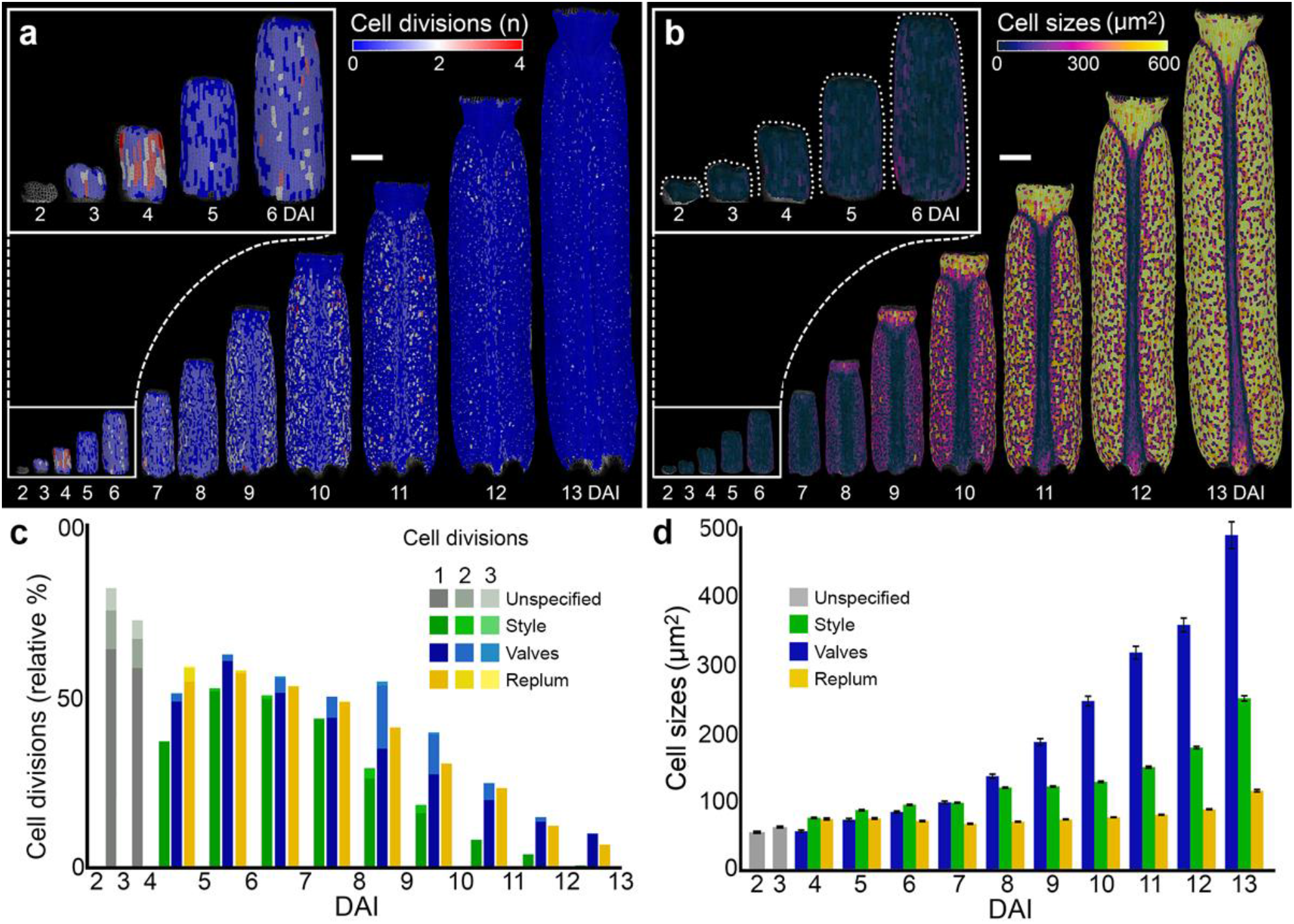
Cellular patterns underlying gynoecium development. **a-b**, Heat-maps of cell divisions (a) and cell sizes (b) for the *Arabidopsis thaliana* gynoecium. **c-d**, Quantifications of cell divisions (c) and cell sizes (d) in different regions of the developing gynoecium. Dotted lines indicate gynoecium outlines. Three independent time lapse series, n=104-10653 cells in one sample at specific time point. Error bars indicate SE. DAI indicates days after gynoecium initiation. Scale bars, 100 µm. Related to Fig. 1

**Extended Data Fig. 2.**
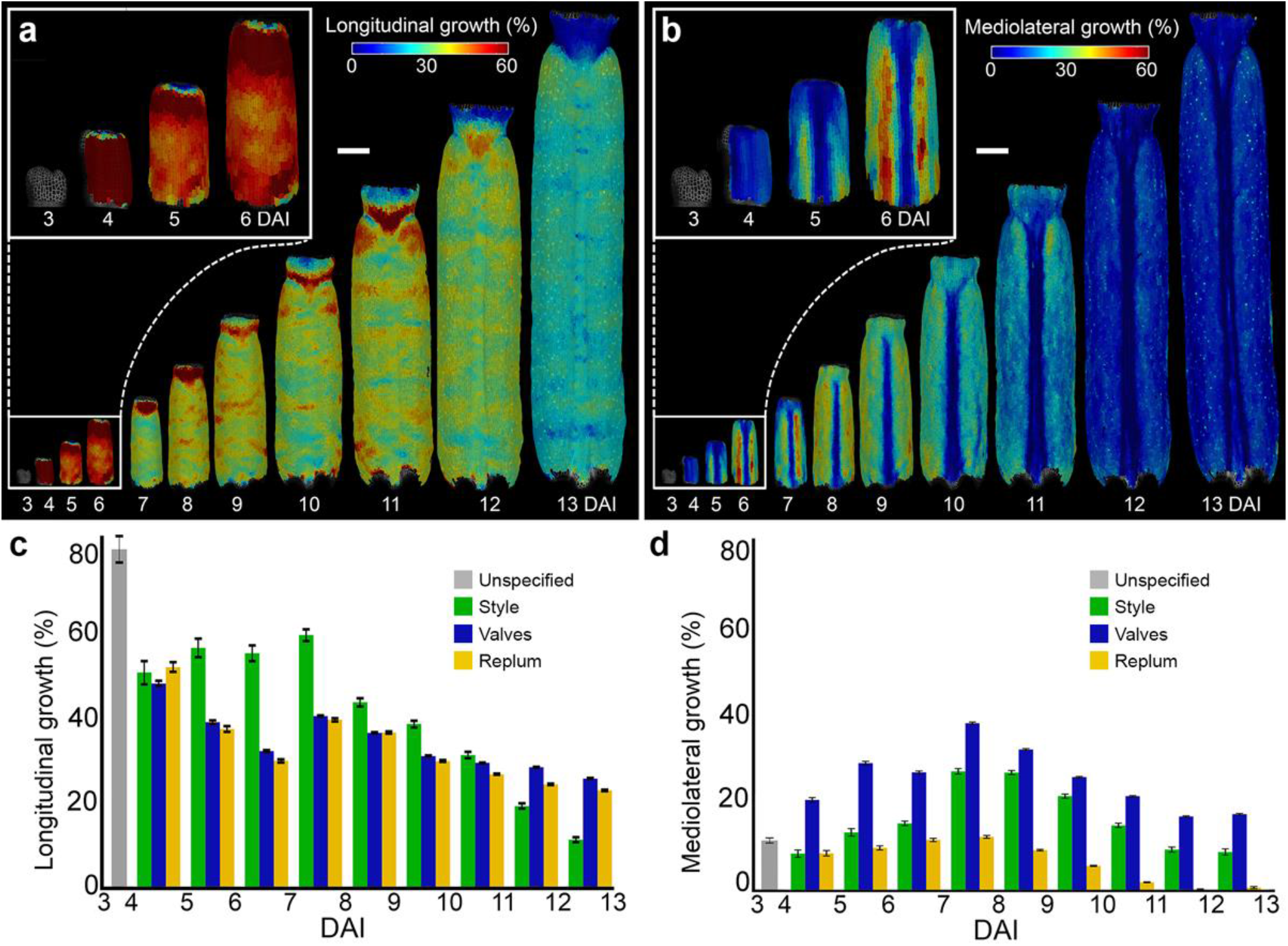
Cellular growth varies along different gynoecium axes. **a-b**, Heat-maps of cellular growth along longitudinal (a) and mediolateral (b) axes of the gynoecium in *Arabidopsis thaliana*. **c-d**, Quantification of cellular growth along longitudinal (**c**) and mediolateral (d) axes of the developing gynoecium. Three independent time lapse series, n=104-10653 cells in one sample at specific time point. Error bars indicate SE. DAI indicates days after gynoecium initiation. Scale bars, 100 µm. Related to Fig. 2.

**Extended Data Fig. 3.**
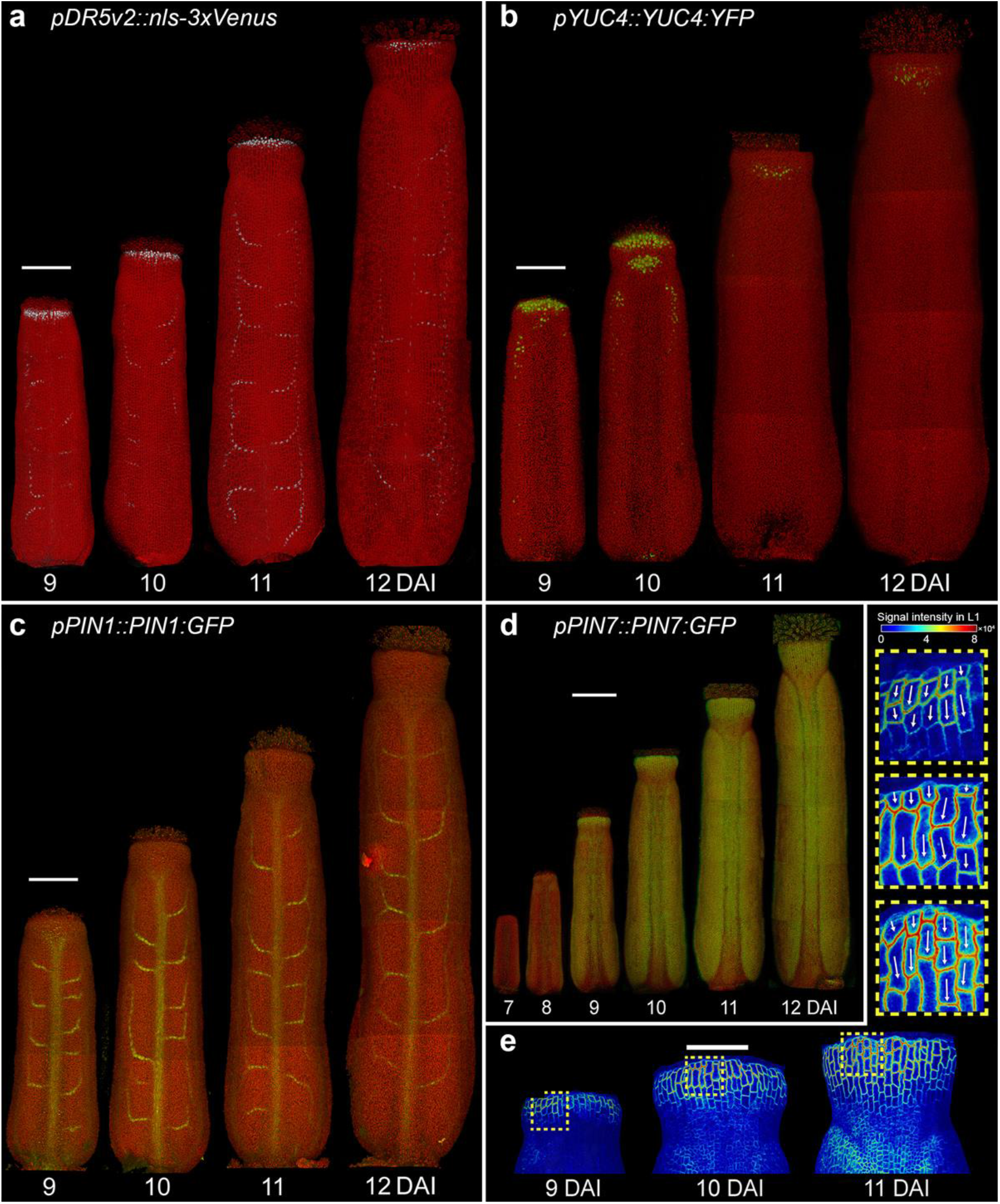
Auxin patterning during gynoecium development. **a-d**, Expression patterns of *pDR5v2::nls-3xVenus* (a), *pYUC4::YUC4:YFP* (b), p*PIN1::PIN1:GFP* (c), and *pPIN7::PIN7:GFP* (d) in the gynoecium of *A. thaliana*. e, Heat-maps of *PIN7:GFP* signal intensity in the L1 epidermal layer of the style. Insets indicate membrane localization; arrows indicate orientation of PIN7 polarization. DAI indicates days after gynoecium initiation. Scale bars, 100 µm (a,b,c,d) and 50 µm (e). Related to Fig. 3.

**Extended Data Fig. 4.**
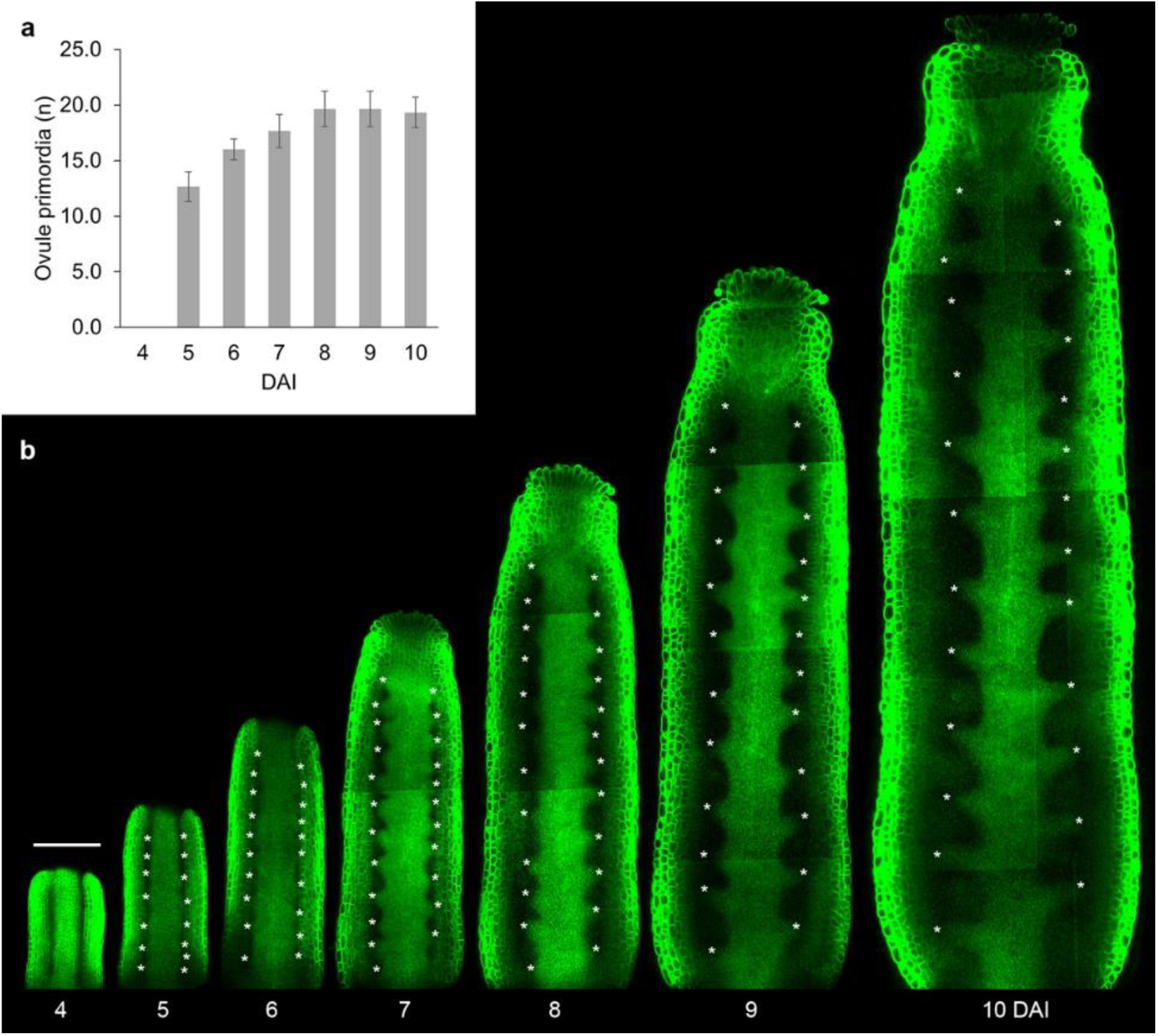
Analysis of the initiation of ovule primordia. **a**, Quantification of ovule primordia in the digital longitudinal sections of the gynoecium of *Arabidopsis thaliana*. Error bars indicate SD (Three independent time lapse series). **b**, Digital longitudinal sections through the gynoecium. Asterisks mark ovule primordia. DAI: days after gynoecium initiation. Scale bar, 100 µm. Related to Fig. 3.

**Extended Data Fig. 5.**
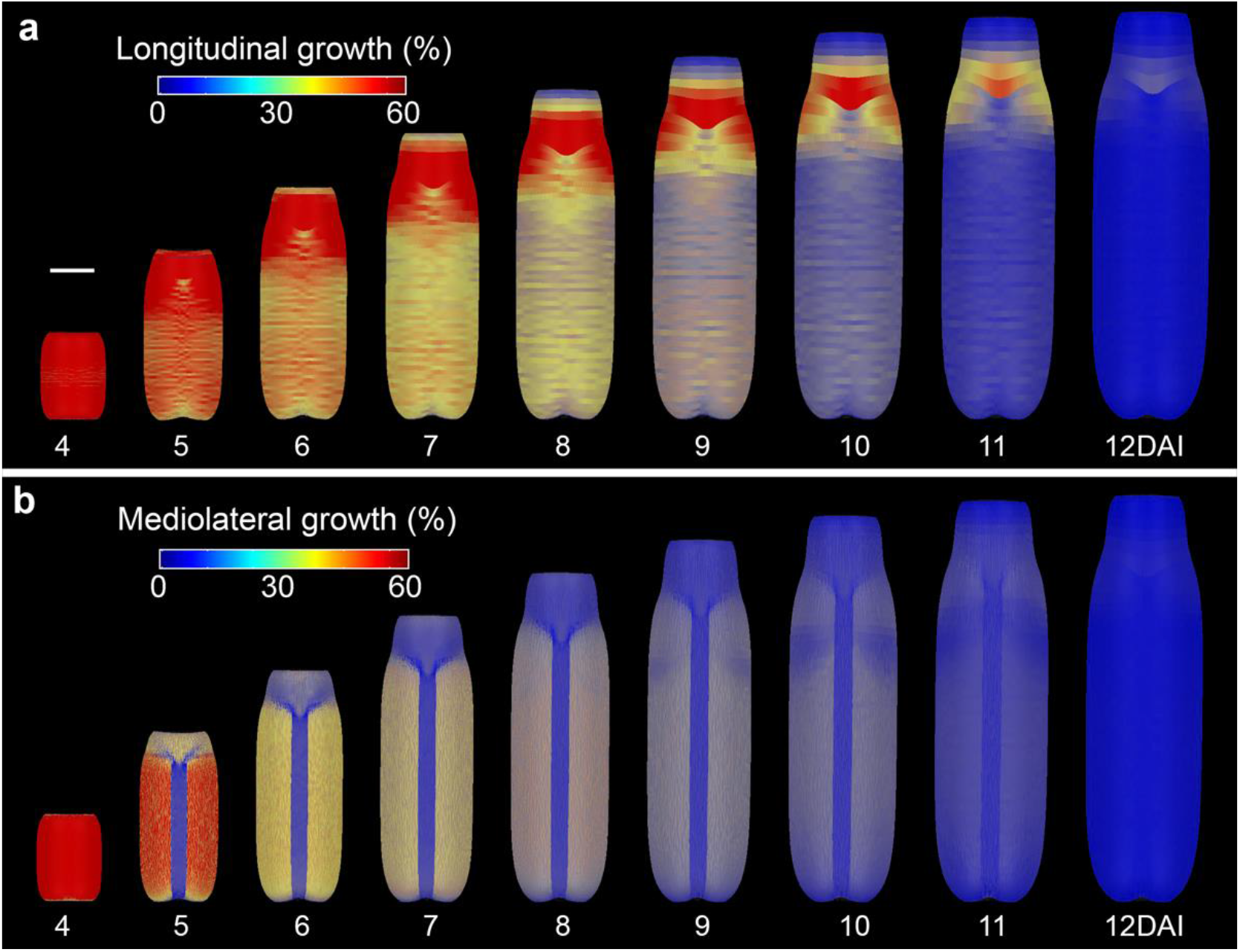
Model of the wild-type gynoecium. **a**, Distribution of longitudinal growth rates predicted by the model of the WT gynoecium development. **b**, Distribution of mediolateral growth rates predicted by the model of the WT gynoecium development. DAI: days after gynoecium initiation. Scale bars, 100 µm. Related to Fig. 3.

**Extended Data Fig. 6.**
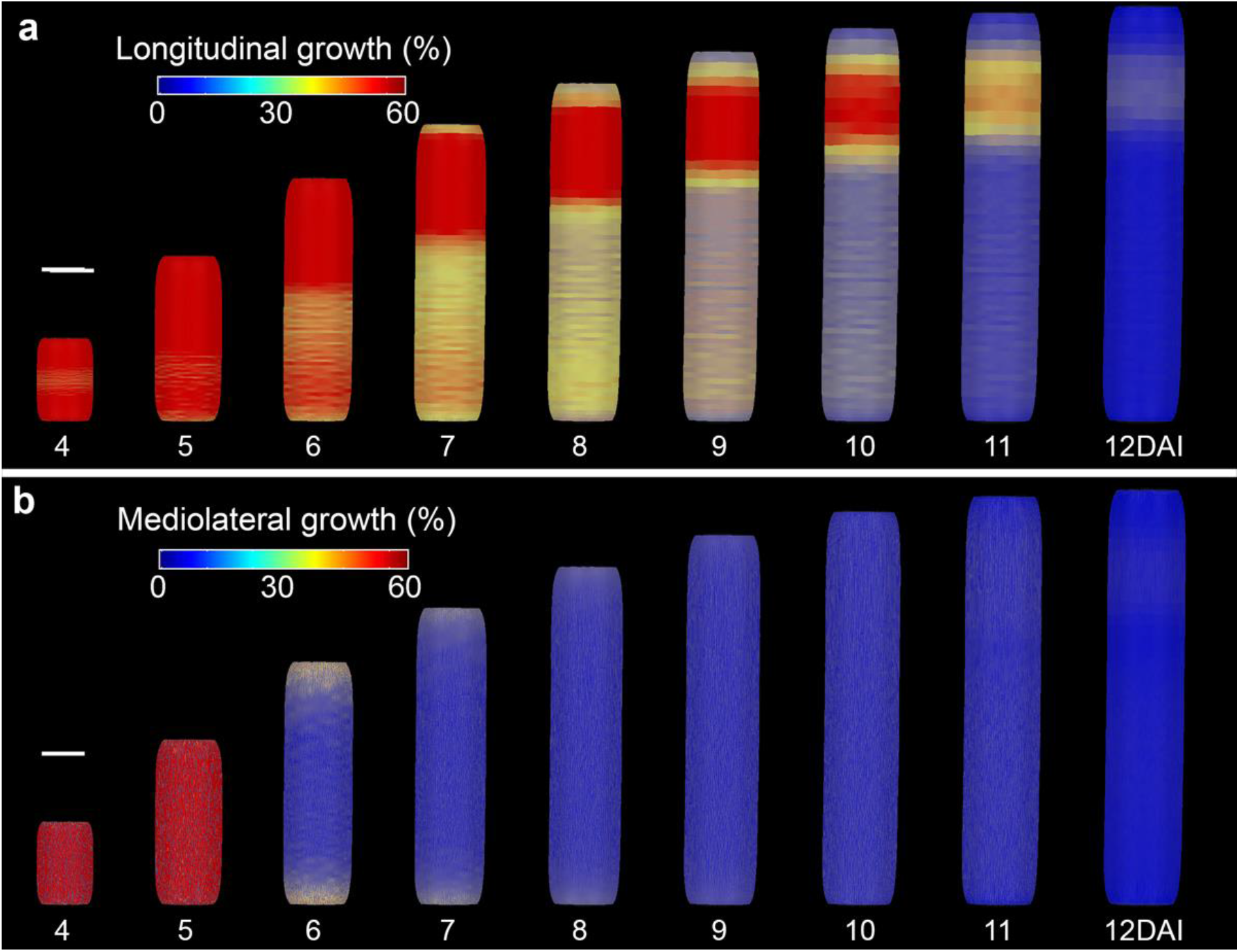
Model of the gynoecium without the CMM. **a**, Distribution of longitudinal growth rates predicted by the model of the NPA-treated gynoecium development. **b**, Distribution of mediolateral growth rates predicted by the model of the NPA-treated gynoecium development. DAI: days after gynoecium initiation. Scale bars, 100 µm. Related to Fig. 4.

**Extended Data Fig. 7.**
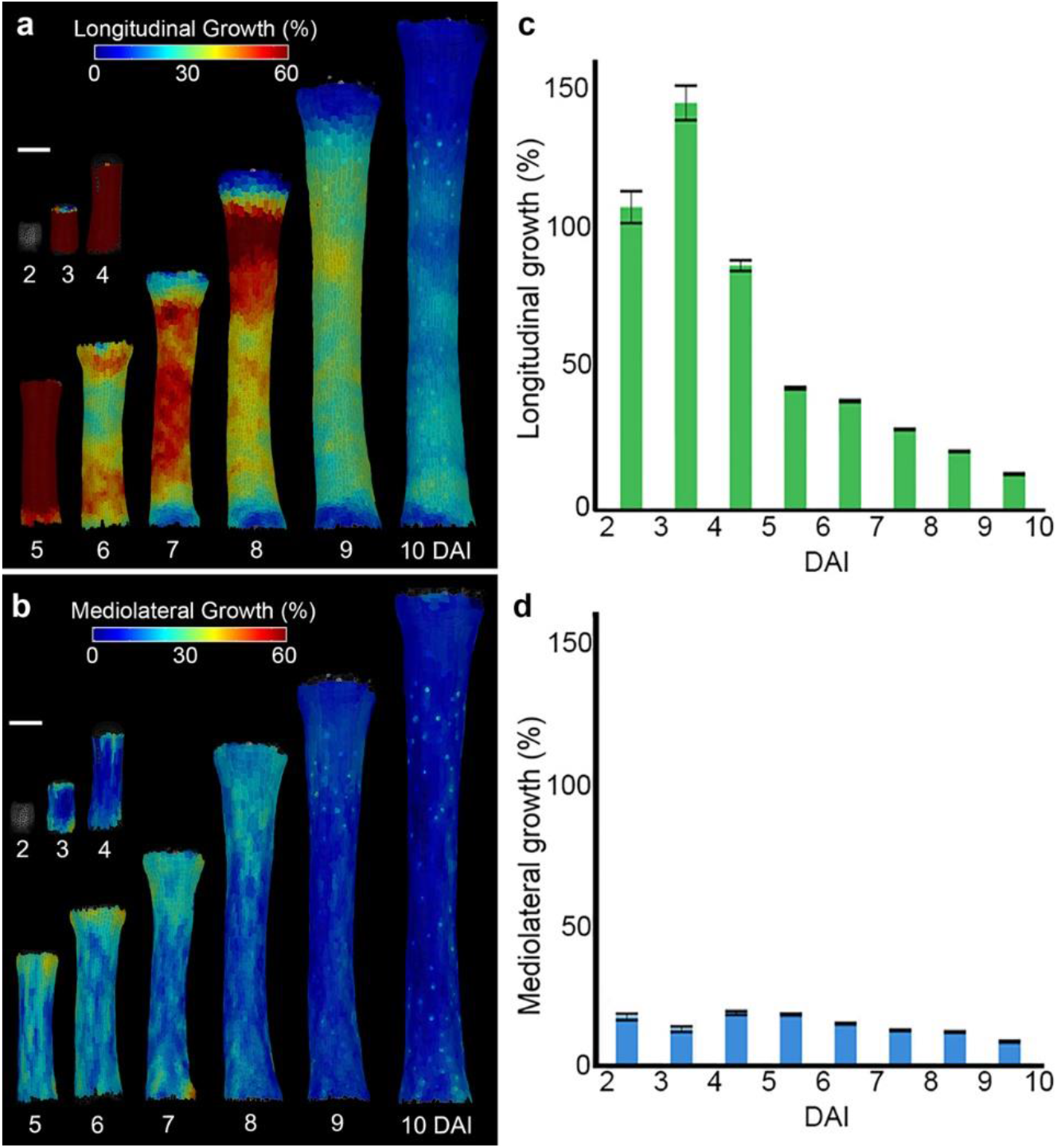
Longitudinal and mediolateral growth in NPA treated gynoecia. **a-b**, Heat maps of longitudinal (a) and mediolateral (b) average growth. **c-d**, Quantification of longitudinal (c) and mediolateral (d) growth in the gynoecia after NPA treatment. Three independent time-lapse series, n=219-1479 cells. DAI: days after gynoecium initiation. Scale bars, 100 µm.

**Extended Data Fig. 8.**
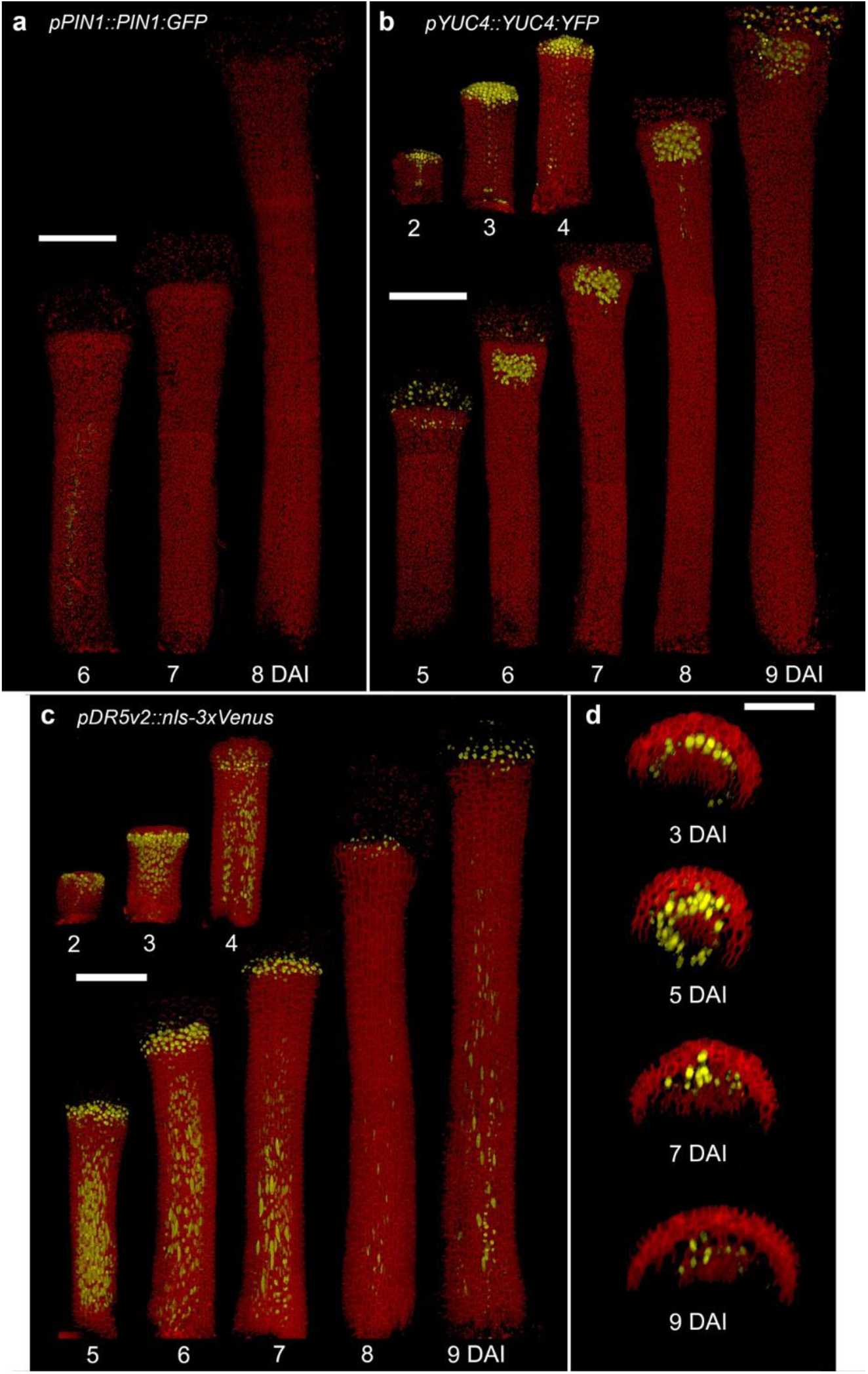
Auxin patterning during the development of valves gynoecium after NPA treatment. **a**, Expression patterns of *pPIN1::PIN1:GFP*. **b**, Expression pattern of *pYUC4::YUC4:GFP*. **c-d**, Expression pattern of *pDR5v2::nls-3xVenus*. DAI: days after gynoecium initiation. Scale bars, 100 µm in (a-c) and 50 µm in (d). Related to Fig. 3.

**Extended Data Fig. 9.**
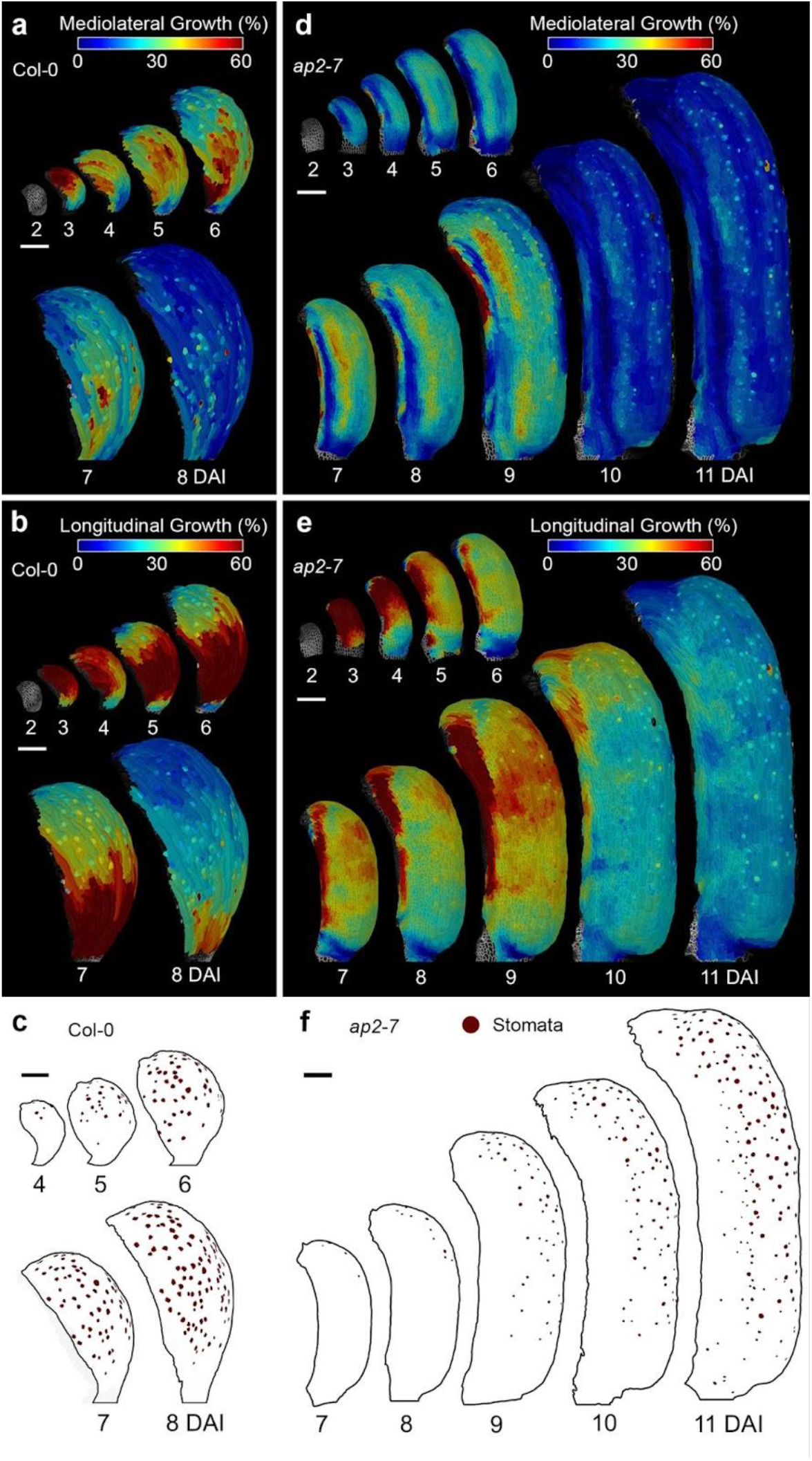
Longitudinal growth, mediolateral growth and stomata distribution in wild-type and *ap2-7* sepals. **a-b**, Heat maps of mediolateral (a) and longitudinal (b) average growth in WT sepals. **c**, Stomata distribution in WT sepal. **d-e**, Heat maps of mediolateral (d) and longitudinal **e**, average growth in *ap2-7* sepals. **f**, Stomata distribution in *ap2-7* sepal. Three independent time lapse series were acquired for wild-type sepals (n=113-1142 cells) and *ap2-7* sepal (n=44-1478 cells). DAI: days after gynoecium initiation. Scale bars, 100 µm. Related to Fig. 4

## Notes

### Competing Interest Statement

The authors have declared no competing interest.

